## Supplementary Information for "Label-free Identification of Protein Aggregates Using Deep Learning"

### Contents:

**Supplementary Note 1: Data processing pipeline and neural network training**

**Supplementary Figure 1: LINA can be used to successfully identify different kinds of aggregates**

**Supplementary Figure 2: Various test set predictions by the LINA pixel-regression model**

**Supplementary Figure 3: Quantitative validation of LINA**

**Supplementary Figure 4: Validation of LINA using negative controls**

**Supplementary Figure 5: Robustness of the trained LINA models**

**Supplementary Figure 6: Validation of LINA on simulated input images of chimeric, non-circular aggregates**

**Supplementary Figure 7: Generalizability of LINA**

**Supplementary Figure 8: LINA accurately predicts aggregates imaged on a commercial microscope**

**Supplementary Figure 9: LINA accurately predicts aggregates imaged using a lower magnification (20X) and NA (0.85) objective**

**Supplementary Figure 10: Morphological distributions of the aggregates in our dataset**

**Supplementary Figure 11: Discrepancy from the complete dataset's standard deviations for different sampling sizes**

**Supplementary Figure 12: Quantitative performance of models trained on different amounts of the training and validation sets**

**Supplementary Figure 13: Dry mass extraction workflow**

**Supplementary Figure 14: Schematic of the microscope setup used in this work**

**Supplementary Figure 15: Schematic of the neural network architecture used in this work (U-Net)**

### Supplementary Note 1: Data processing pipeline and neural network training

We used custom MATLAB (R2021a) (Mathworks) scripts (available here) to retrieve the phase information from the brightfield images and produce quantitative phase images. These scripts are also used for pixel-registration in the 8 z-planes for both phase and fluorescence images. The images are cropped to a size of 352 pixels x 352 pixels.

We used Fiji (v2.9.0) scripts to produce maximum z-projection fluorescence images, to segment these images using Otsu thresholding and produce the labels for pixel classification, to prepare the color-coded maximum z-projection phase image shown in Fig. 2a, and for the image processing in Fig. 2b. Fiji was also used to segment network predictions and produce masks which are used to measure the area, circularity or dry mass of aggregates. Supplementary Fig. 13 summarizes the dry mass extraction process.

Our models are trained on a deep CNN with a U-Net[1] architecture (Supplementary Fig. 15). Compared to the original architecture, we reduced the number of feature maps by a factor of 4 which led to a reduction in the trainable parameters by a factor of 15. This notably reduces GPU memory usage and training time, while still enabling excellent performance. We used TensorFlow (v2.8.0) and Keras (v2.8.0) to build our network, and training was done on a workstation equipped with an NVIDIA GeForce RTX 3090 GPU. The input images have dimensions of 352 pixels x 352 pixels in XY as aforementioned. Some models are trained on 8-plane inputs and some use fewer planes (4, 2, or 1). We used the adaptive moment estimation (Adam) optimizer with a learning rate of 1e-4 and a mean squared error loss function. Before training our models, we normalize both the phase and fluorescence images by rescaling each image to be between 0 and 1. We used 10% of our dataset as the test set and split the rest of the dataset into training and validation sets with 20% being used for validation. We used the 'EarlyStopping' (on the validation loss) and 'ModelCheckpoint' callbacks to avoid overfitting and to save the best, most general models.

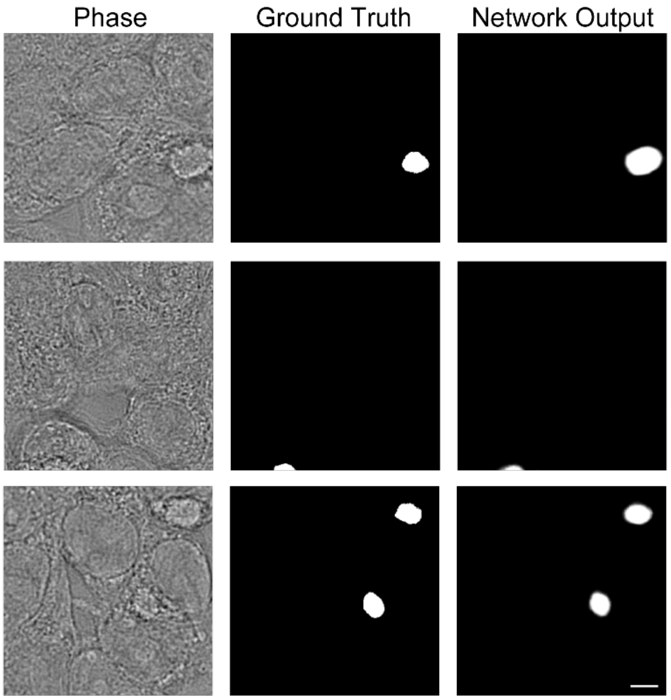


**Supplementary Figure 1: LINA can be used to successfully identify different kinds of aggregates.** The performance of the pixel classification model is shown for three different test set examples: one aggregate, an aggregate on the edge of the frame, and two aggregates in one frame. Scale bar: 5 μm.

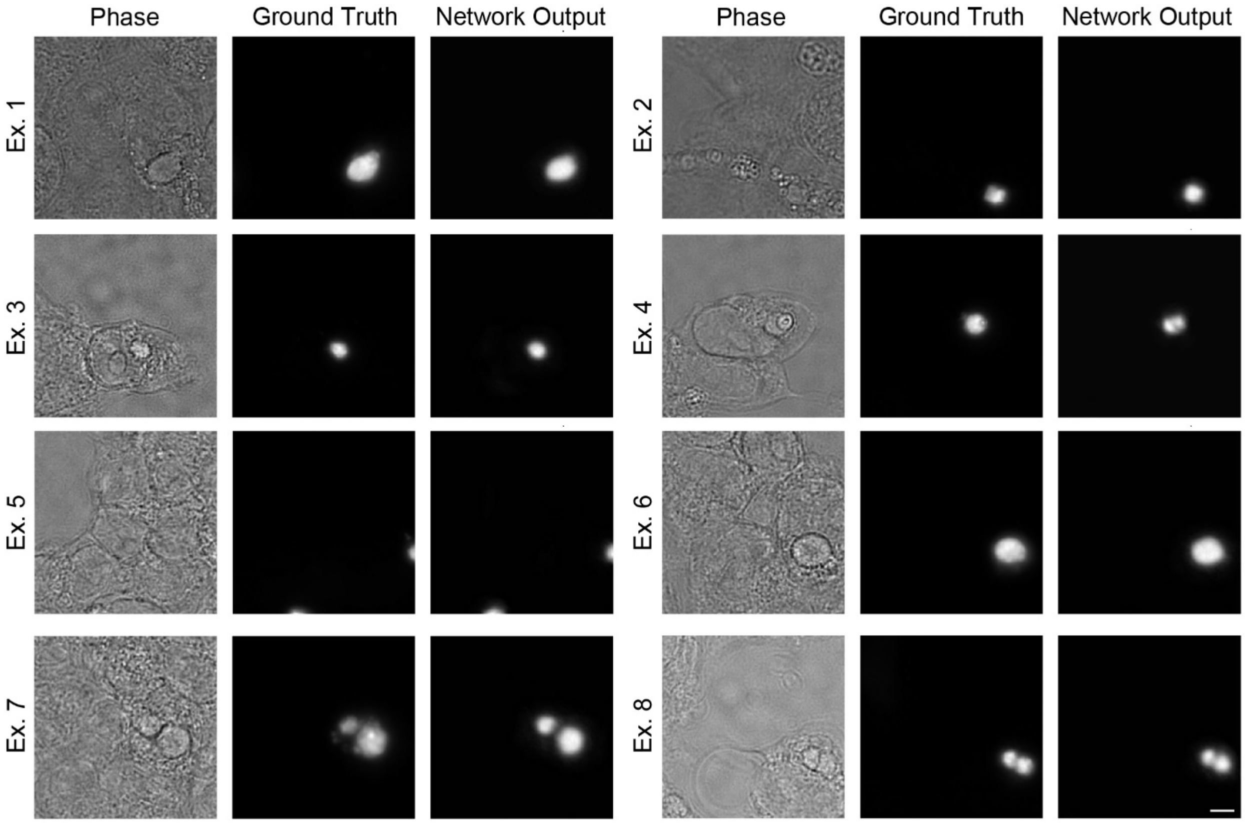


**Supplementary Figure 2: Various test set predictions by the LINA pixel-regression model.** The model performs consistently well over a range of examples, including aggregates on the edges of the frame and aggregates that are spatially near each other. Scale bar: 5 μm.

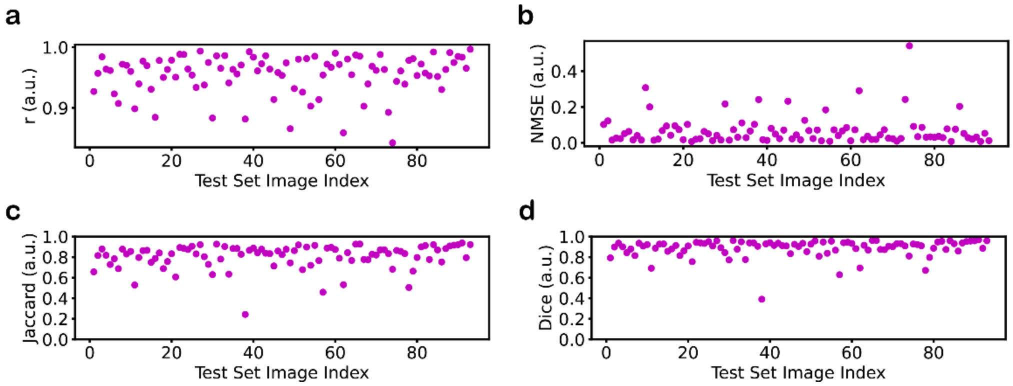


**Supplementary Figure 3: Quantitative validation of LINA.** (a) Pearson correlation coefficient (r) computed only on the regions where there are aggregates. (b) Normalized mean squared error computed only on the regions where there are aggregates. (c) Jaccard index. (d) Dice coefficient.

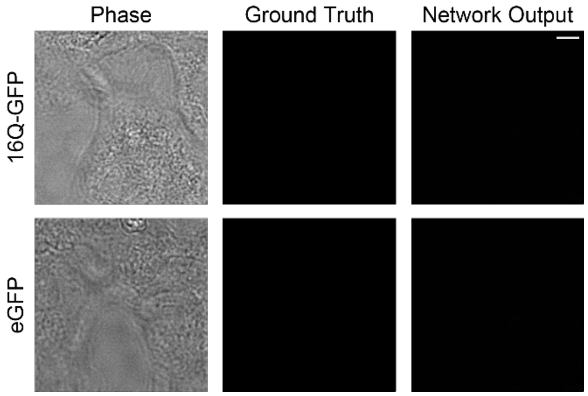


**Supplementary Figure 4: Validation of LINA using negative controls.** (a) Example images visualizing the pixel-regression model's performance on a negative control, 16Q-GFP. Protein aggregates are not formed at this polyQ length. (b) Example images visualizing the pixel-regression model's performance on a negative control, eGFP, where there is no Httex1 protein at all. In both (a) and (b) the model successfully predicts the absence of aggregates. Scale bar: 5 μm.

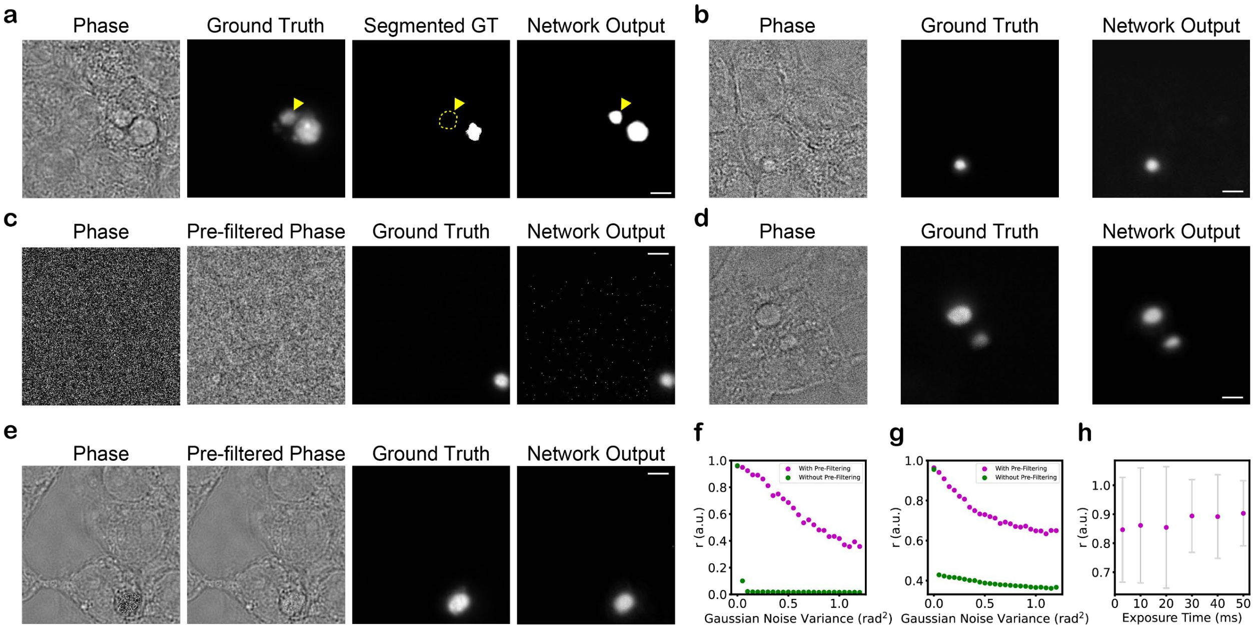


**Supplementary Figure 5: Robustness of the trained LINA models.** (a) The pixel classification network can outperform the label and correctly identify unsegmented aggregates. A test set example is shown in which an aggregate was correctly identified by the neural network, despite being missing in the segmented ground truth label due to a segmentation error. The fluorescence ground truth signal confirms the presence of an aggregate in those pixels. (b), (c), (e), (f), and (g) show the pixel-regression model's behavior when noise is artificially added. (b) The model is still able to identify the aggregate in a test set example in which a small amount of Gaussian noise is added. (c) Images of a test set example at an extreme level of artificially added noise (0.48 rad$^2$ variance). The model is still able to identify the aggregate after the input is pre-filtered, despite both the original and pre-filtered phase images looking like pure noise. (f) At larger amounts of artificially added noise, the model performs worse quantitatively, however, this is mitigated by pre-filtering (blurring) the images prior to them being given as input to the network. With pre-filtering, the model remains stable even at extremely large (unrealistic) amounts of noise, as shown in (c). (e) Images of a test set example where noise (0.2 rad$^2$ variance) is artificially added to the region where there is an aggregate. The model is still able to identify the aggregate after the input is pre-filtered. (g) Similar to (f), the quantitative performance is maintained after pre-filtering but degrades slowly until it saturates. (d) and (h) illustrate LINA's robustness at varying signal-to-noise ratios. (d) With an image acquired at a 3 ms exposure time, LINA is still able to identify aggregates, even though it was trained exclusively using images taken at a 50ms exposure time. (h) Summary of LINA's quantitative performance for input images taken at different exposure times, showing high accuracy and consistency. Error bars represent the standard deviation. Scale bars: 5 μm.

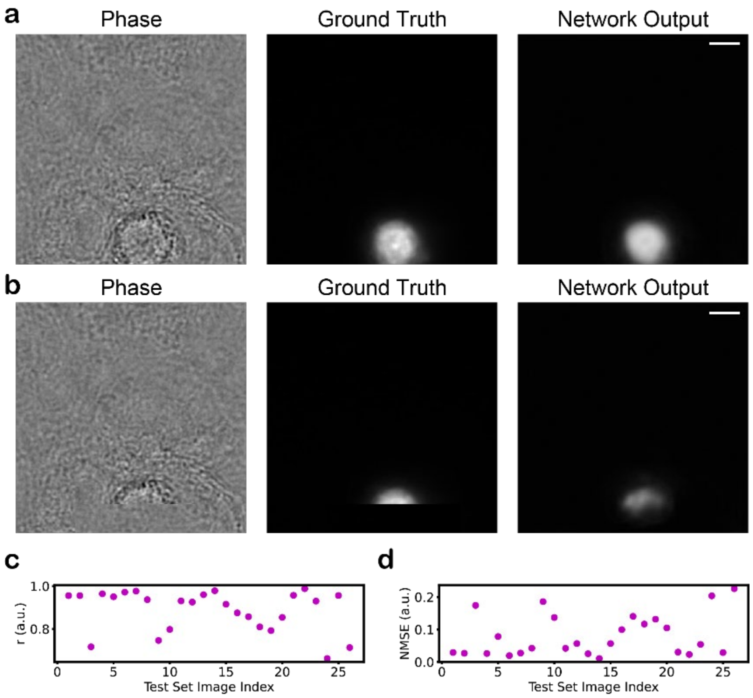


**Supplementary Figure 6: Validation of LINA on simulated input images of chimeric, non-circular aggregates.** (a) Example images visualizing the model's performance on a non-simulated, circular aggregate. (b) The model's performance on the same example after replacing part of the aggregate in the input image and the ground truth with part of a cell/background image (and its corresponding fluorescence signal) to create a simulated test example of a chimeric, non-circular aggregate. The model is able to accurately predict the presence of the heterogenous aggregate. (c) Pearson correlation coefficient (r) computed on the simulated test set, only on the regions where there are aggregates. The metric is computed for the 8-plane-QPI pixel-regression model. (d) Normalized mean squared error computed on the simulated test set, only on the regions where there are aggregates. The metric is computed for the 8-plane-QPI pixel-regression model. Scale bars: 5 μm.

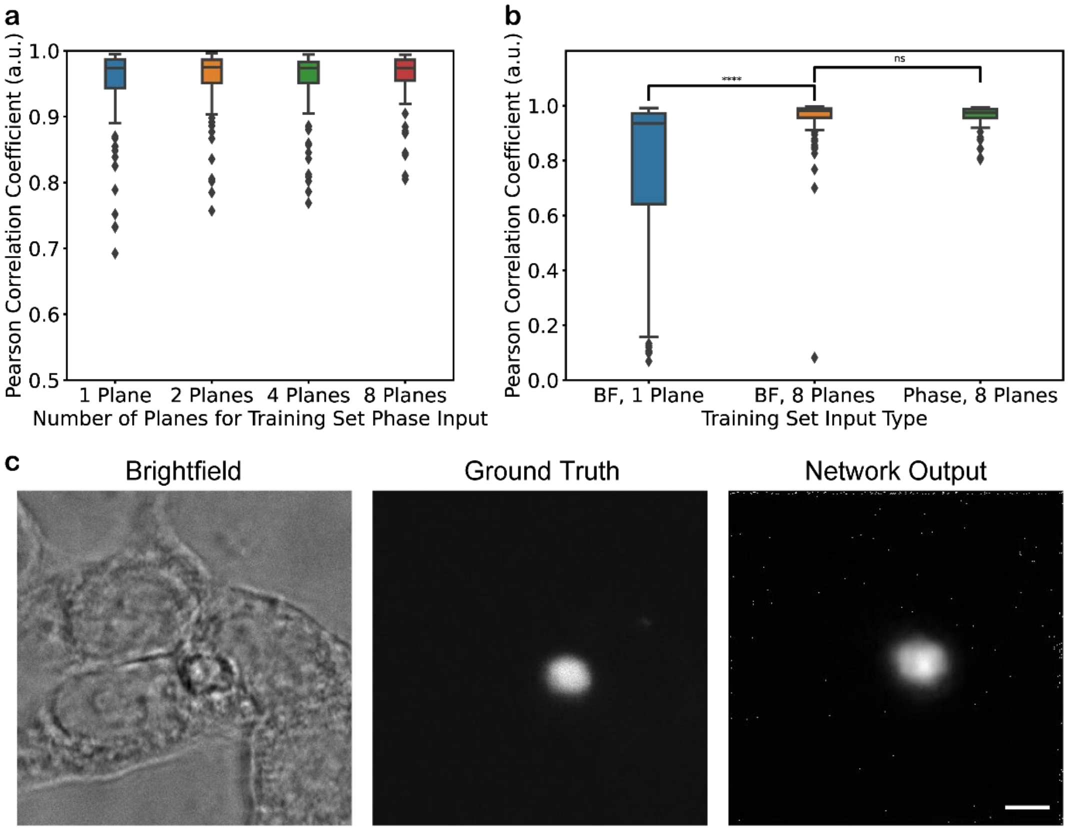


**Supplementary Figure 7: Generalizability of LINA.** (a) Quantitative performance of the pixel-regression model when trained by QPI inputs with different numbers of planes. (b) Quantitative performance of the pixel-regression model when trained by brightfield inputs with one or 8 planes, compared with the model trained on 8-plane QPI inputs. (a, b) Unpaired, two-sided t-tests were performed. ns: $p > 0.05$, ****: $p \leq 0.0001$. The box plots show the median (central line), first and third quartile (upper and lower lines), 1.5x the interquartile range (whiskers), and the outliers (markers). (c) Example of an image acquired on a different imaging setup, with the aggregate being identified by the 1-plane brightfield model. Scale bar: 5 μm.

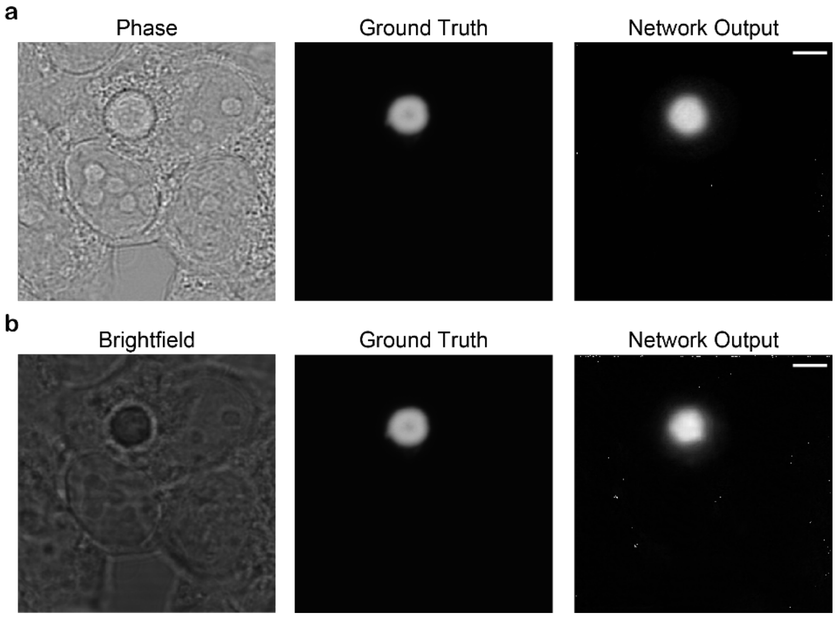


**Supplementary Figure 8: LINA accurately predicts aggregates imaged on a commercial microscope.** (a) Example images showing LINA's performance on an 8-plane QPI input. (b) Example images showing LINA's performance on a 1-plane brightfield input. Scale bars: 5 µm.

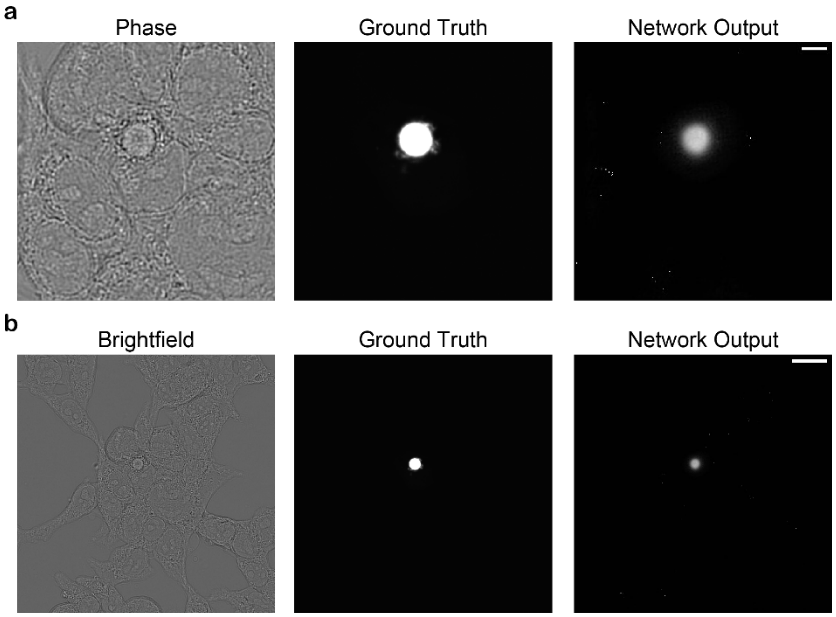


**Supplementary Figure 9: LINA accurately predicts aggregates imaged using a lower magnification (20X) and NA (0.85) objective.** (a) Example images showing LINA's performance on an 8-plane QPI input, which is cropped around the aggregate to have the standard input size of 352 pixels x 352 pixels. Scale bar: 5 µm. (b) Example images showing LINA's performance on the 1024 pixels x 1024 pixels input directly. Scale bar: 20 µm.

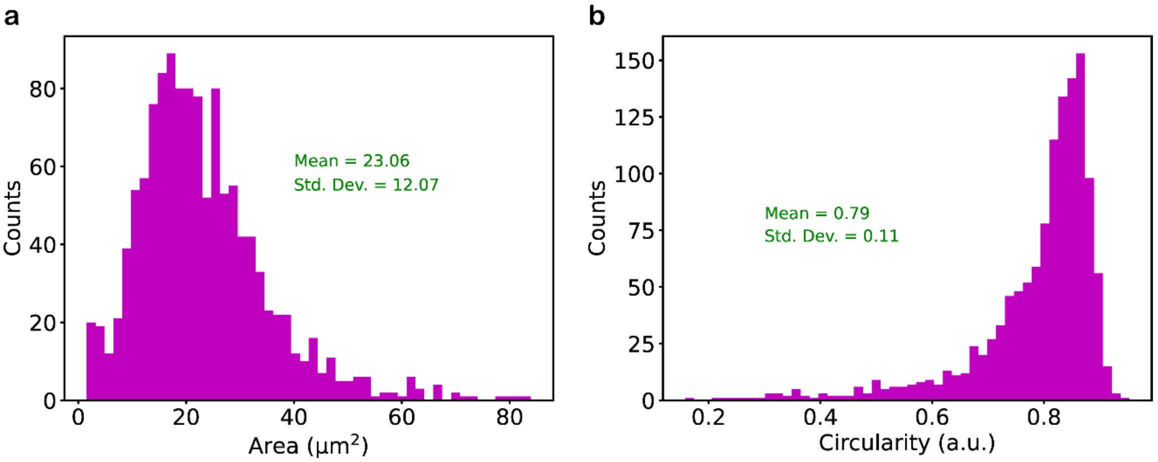


**Supplementary Figure 10: Morphological distributions of the Httex1-72Q-GFP aggregates in our dataset.** (a) Histogram showing the area distribution. (b) Histogram showing the circularity distribution.

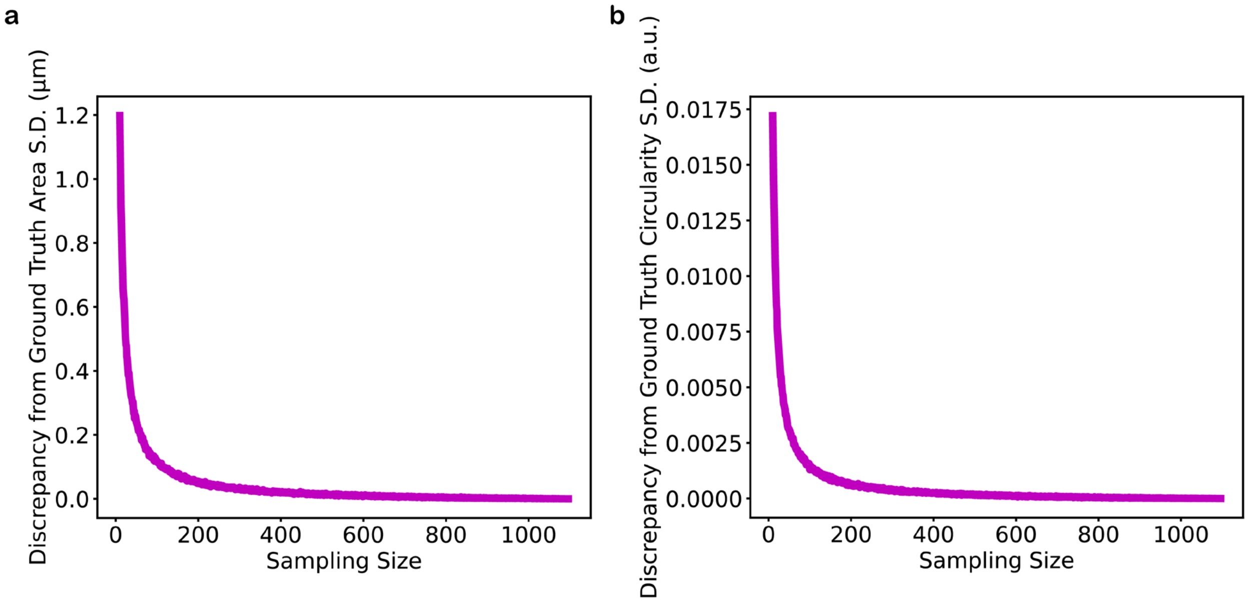


**Supplementary Figure 11: Discrepancy from the complete dataset's area and circularity standard deviations for different sampling sizes.** This simulation is used to estimate how much data is needed to capture the diversity of our complete dataset, before reaching diminishing returns. At each sampling size, random sampling is repeated 100,000 times.

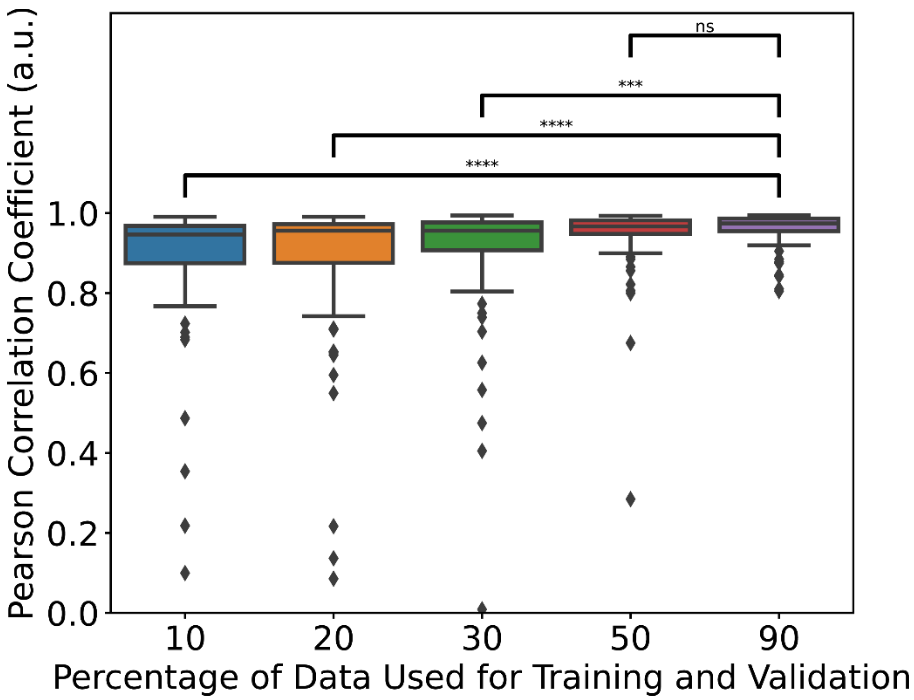


**Supplementary Figure 12: Quantitative performance of models trained on different amounts of the training and validation sets.** Unpaired, two-sided t-tests were performed. ns: p > 0.05, **: p ≤ 0.01, ***: p ≤ 0.001, ****: p ≤ 0.0001. The box plot shows the median (central line), first and third quartile (upper and lower lines), 1.5x the interquartile range (whiskers), and the outliers (markers).

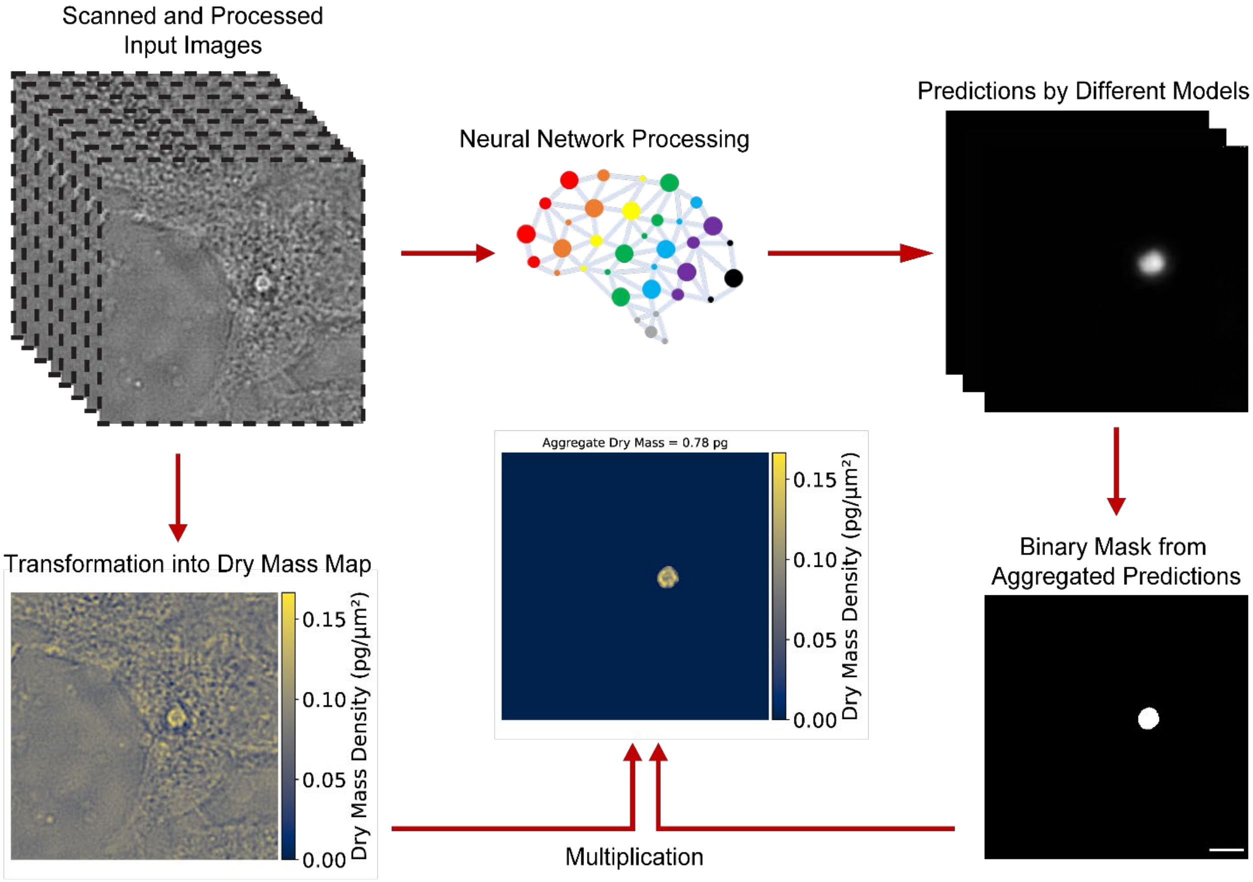


**Supplementary Figure 13: Dry mass extraction workflow.** Images are acquired at various FOVs for each protein construct using an automatic-scanning method. The images are then processed and used as input to neural network models. The models identify FOVs which have aggregates and localize them. For each FOV with an aggregate, a binary mask is then generated from the aggregated predictions of these models, and it is multiplied by a dry mass map which is generated from the QPI image, yielding a dry mass map solely of the identified aggregate at each FOV. By integrating over the area of the aggregate in the dry mass map, it is possible to measure its dry mass. Scale bar: 5 μm.

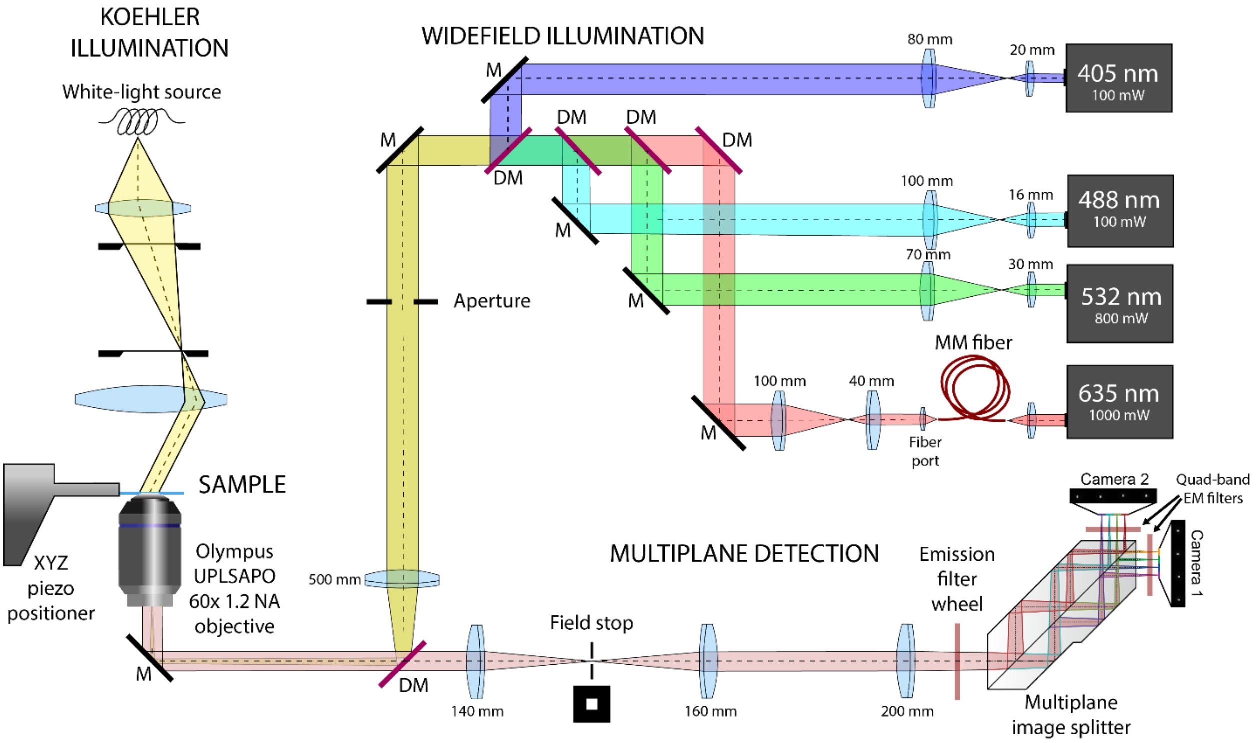


**Supplementary Figure 14: Schematic of the microscope setup used in this work.** M – mirror, DM – dichroic mirror, MM – multi-mode, EM – emission. Adapted from work by Navikas and colleagues[4]. First developed and described in work by Descloux and colleagues[5].

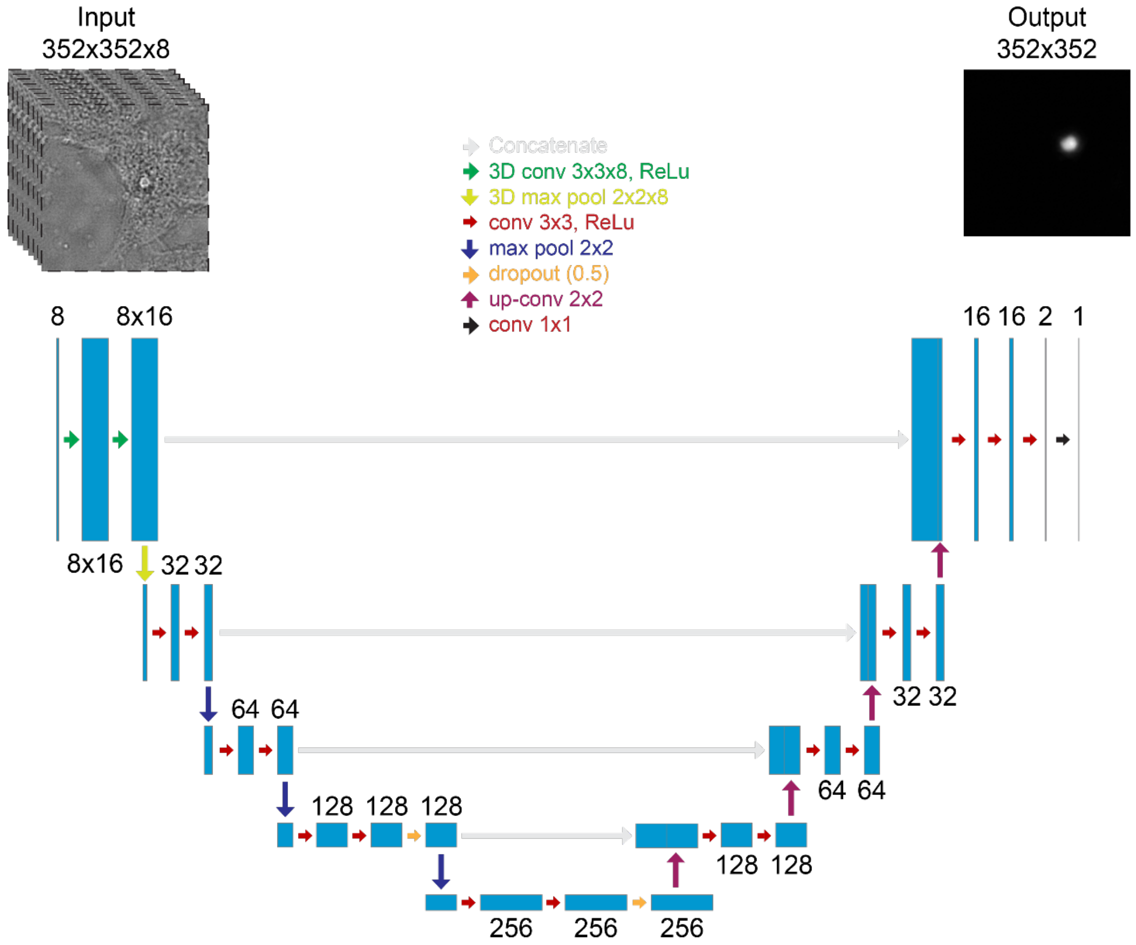


**Supplementary Figure 15: Schematic of the neural network architecture used in this work (U-Net).** Note that we also train models with fewer-plane inputs. Adapted from work by Ronneberger and colleagues[1] and work from Ounkomol and colleagues[6].
